# Arginase Inhibition Supports Survival and Differentiation of Neuronal Precursors in Adult Alzheimer’s Disease Mice

**DOI:** 10.1101/2019.12.26.888727

**Authors:** Baruh Polis, Vyacheslav Gurevich, Naamah Bloch, Abraham O. Samson

## Abstract

Adult neurogenesis is a complex physiological process, which plays a central role in maintaining cognitive functions, and consists of progenitor cell proliferation, newborn cell migration, and cell maturation. Adult neurogenesis is susceptible to alterations under various physiological and pathological conditions. A substantial decay of neurogenesis has been documented in Alzheimer’s disease (AD) patients and animal AD models; however, several treatment strategies can halt any further decline and even induce neurogenesis.

Our previous results indicated a potential effect of arginase inhibition, with norvaline, on various aspects of neurogenesis in triple-transgenic mice. To better evaluate this effect, we chronically administer an arginase inhibitor, norvaline, to triple-transgenic and wild-type mice, and apply an advanced immunohistochemistry approach with several biomarkers and bright-field microscopy.

Remarkably, we evidence a significant reduction in the density of neuronal progenitors, which demonstrate a different phenotype in the hippocampi of triple-transgenic mice as compared to wild-type animals. However, norvaline shows no significant effect upon the progenitor cell number and constitution. We demonstrate that norvaline treatment leads to an escalation of the polysialylated neuronal cell adhesion molecule immunopositivity, which suggests an improvement in the newborn neuron survival rate. Additionally, we identify a significant increase in the hippocampal microtubule-associated protein 2 stain intensity. We also explore the molecular mechanisms underlying the effects of norvaline on adult mice neurogenesis and provide insights into their machinery.

## Introduction

The adult murine brain continuously generates neuronal progenitor cells (NPCs) in the subventricular zone (SVZ) and subgranular zone (SGZ) of the hippocampal dentate gyrus [1]. However, the existence of human adult neurogenesis has been a subject of intense scientific debate, until recently. Tobin *et al*. (2019) evidenced hippocampal neurogenesis persisting throughout life in the brains of centenarians and even of Alzheimer’s disease (AD) patients [2]. Convincingly, they demonstrated that the density of NPCs, neuroblasts, and immature neurons significantly decreases in cases of mild cognitive impairment and in clinical AD as compared to healthy controls.

In addition, various animal models of AD are characterized by diminished adult neurogenesis. In particular, triple-transgenic mice (3×Tg) show an age-dependent neurogenesis insufficiency that is detectable in the hippocampus starting at four months of age [3]. Of note, the decline of neurogenesis in this AD model precedes the manifestation of classical hallmarks of AD pathology, such as deposition of amyloid plaques and neurofibrillary tangles in the brain, as well as memory impairment. Remarkably, as a form of neuroplasticity, adult neurogenesis has been shown to modulate vulnerability to degenerative processes and influence the course of AD [4]. Moreover, various supporting adult neurogenesis treatment strategies have demonstrated their competence to counteract the pathological behavioral outcomes in murine AD models [5].

Nevertheless, the mechanisms that regulate NPC proliferation, differentiation, and migration remain poorly understood. Several reports have demonstrated the effects of nitric oxide (NO) donors upon the rate of NPC proliferation and migration. Zhang *et al*. (2001) evidenced a substantial increase in the rate of SGZ neurogenesis following treatment with NO donors in rats [6]. Lu *et al*. (2003) demonstrated upregulation of neurogenesis and a reduction in functional deficits following the administration of an NO donor after traumatic brain injury in rats. [7]. Chen *et al*. (2004) proved the potency of a complex therapy for stroke with an NO donor and human bone marrow stromal cells to enhance neurogenesis in rats [8]. In contrast, nitric oxide synthase (NOS) deficiency has been shown to impede neurogenesis. Reif *et al*. (2004) reported a significant decrease in NPC proliferation in the SGZ of NOS3 knockout mice, accompanied by a decline in the levels of vascular endothelial growth factor (VEGF) [9]. These authors suggest a principal role of NOS3 in the stimulation of neurogenesis. Other groups have shown that the effects of NOS3 on progenitor cells are mediated via VEGF. Chen *et al*. (2005) showed that NOS3 is a downstream mediator of VEGF [10]. They suggest that NOS3 regulates brain-derived neurotrophic factor (BDNF) expression in the ischemic brain and influences NPC proliferation and migration. A more recent report by Jin *et al*. (2017) revealed that endogenous neuronal NOS1 positively regulates neurogenesis [11]. These authors demonstrated that NOS1 repression decreases neuronal differentiation, and *vice versa*, NOS1 upregulation promotes it.

The semi-essential amino acid, arginine, is a mutual substrate for both NOS and arginase. Brain arginine deprivation, due to arginase over-activation, has been suggested as a cause of AD [12]. Consequently, arginine supplementation [13] and/or arginase inhibition have been proposed to halt AD development [14], and have been successfully tested in AD mice [15]. Previously, we showed that arginase inhibition with norvaline upsurges the hippocampal levels of NOS3 [16], and NOS1 [17] in AD model mice.

Additionally, our advanced proteomics assay revealed that chronic treatment of 3×Tg mice with norvaline led to the activation of several critical for adult neurogenesis biological processes [15]. One of the most significant pathways detected was the neuregulin (NRG) pathway. Of note, NRGs comprise a cluster of epidermal growth factor-like proteins that are highly involved in neural development and brain homeostasis [18]. Accumulating evidence suggests a strong impact of NRG1 signaling upon cognitive function and neuropathology in AD. The overexpression of NRG1 in the hippocampus of AD mice improves memory and ameliorates disease-associated neuropathology [19]. Moreover, the systemic administration of NRG1 intensifies neurogenesis in the mouse dentate gyrus [20-21].

Previously, we demonstrated that norvaline treatment up-regulated the VEGF signaling pathway in 3×Tg mice [15]. We also noted that NOS3 mediates VEGF activity [10]. Since norvaline elevates NOS3 levels [16], we suggest a significant involvement of VEGF activation in the phenotype observed following norvaline treatment. VEGF is essential for neuroprotection [22], and its activation is beneficial for individuals with early signs of AD-associated dementia [23]. VEGF preconditioning has been shown to attenuate age-related decay of adult hippocampal neurogenesis in mice [24]. Additionally, the administration of VEGF in rats with focal cerebral ischemia reduces the infarct size and enhances neurogenesis [25].

Likewise, we disclosed that norvaline treatment led to a significant increase in the levels of the glial cell-derived neurotrophic factor (GDNF) receptor RET (REarranged during Transfection) [15], which is a common signaling receptor for GDNF-family ligands [26]. Of note, GDNF is down-regulated in 3×Tg mice [27], and its overexpression improves cognitive function in this AD model [28]. Moreover, GDNF supports neuronal survival [29], and RET is essential for mediating the neuroprotective and neuroregenerative effects of GDNF [30]. This factor has been shown to increase neurogenesis in the adult hippocampus [31]. Furthermore, neural cell adhesion molecule (NCAM), a second signaling receptor of GDNF [32], demonstrated a 43% level elevation following norvaline treatment [15]. Of note, NCAM regulates synaptic plasticity [33] and mediates axonal growth in hippocampal and cortical neurons [32]. Consequently, an upsurge in its hippocampal levels points to the treatment-associated improvement of neuroprotective mechanisms in 3×Tg mice brains.

Growing evidence indicates that another neuroprotective factor, nerve growth factor (NGF), plays a crucial role in the pathogenesis of AD [34]. NGF administration was shown to promote neurogenesis in adult rodents [35]. Accordingly, NGF application has emerged as a promising approach in AD therapy [36]. The neuroprotective effects of NGF are mediated via tropomyosin receptor kinase A [37], which demonstrated a significant 56% increase following norvaline treatment [15].

Additionally, we disclosed a significant, more than two-fold elevation, in the levels of neuroligin-1 in the hippocampi of 3×Tg mice following treatment with norvaline [16]. Neuroligin-1 knockdown has been shown to reduce the survival rate of adult-generated newborn hippocampal neurons [38]. This group previously demonstrated that neuroligin-1 overexpression selectively increases the degree of excitatory synapse formation in adult mice [39].

Overall, several lines of converging evidence unequivocally point to the manifold effects of norvaline upon neurogenesis, neuronal cell differentiation, and migration. Consequently, it is plausible to hypothesize that this substance induces adult neurogenesis in AD model mice. In order to check this hypothesis, we applied an innovative immunohistochemistry approach, using several bio-markers and bright-field microscopy, to study the effects of arginase inhibition with norvaline upon the rate of neurogenesis in 3×Tg mice. We demonstrate that norvaline promotes neuronal differentiation and survival. We also explore the molecular mechanisms underlying the effects of norvaline on adult neurogenesis and provide insights into their machinery.

## Methods

### Strains of mice and treatment

Homozygous 3×Tg mice, harboring PS1(M146V), APP(Swe), and tau(P301L) transgenes, were purchased from Jackson Laboratory (Bar Harbor, ME, USA) and bred in our animal facility. These mice exhibit memory deficits associated with amyloid plaques deposition and tangle pathology [40].

Randomly chosen, male 4-month-old transgenic mice and age-matched male C57Bl/6 mice (wild-type) were divided into four groups (10 mice in each group) and housed in individually ventilated cages (Lab Products Inc., Seaford, DE, USA), with five mice per cage. The animals were provided with water and food *ad libitum*. The control animals received regular water. The experimental mice received water with dissolved (250 mg/L) norvaline (Sigma, St. Louis, MO, USA). The experiment lasted 2.5 months. The experiments were performed according to the “Guide for the Care and Use of Laboratory Animals” [41] and the experimental protocol (𝒩º 82-10–2017) was approved by Bar-Ilan University animal care and use committee.

### Tissue preparation and slicing

Four animals from each group were deeply anesthetized, with an intraperitoneal injection of 0.2 ml Pental (CTS Chemical Industries, Kiryat Malachi, Israel). The animals were perfused transcardially with ice-cold phosphate buffer saline (PBS), followed by ice-cold paraformaldehyde 4% in PBS. The mice were decapitated, and their brains were carefully removed and fixed in 4% paraformaldehyde for 24 hours and then were transferred to 70% ethanol at 4°C for 48 hours. The tissues were dehydrated and paraffin-embedded. The paraffin-embedded tissue blocks were chilled on ice and sliced on Leica RM2235 manual rotary microtome to a thickness of four µm. Then, the sections were mounted onto gelatin-coated slides, dried overnight at room temperature, and stored at 4 °C in slide storage boxes.

### Quantitative immunohistochemistry

We studied neurogenesis within the dentate gyrus of the adult mice hippocampal formation by means of immunohistochemistry. Staining was accomplished on the Leica Bond Max system (Leica Biosystems Newcastle Ltd, UK). Brain sections were dewaxed and pretreated with the epitope-retrieval solution (ER, Leica Biosystems Newcastle Ltd, UK), and then incubated for 30 minutes with primary antibodies. A Leica Refine-HRP kit (Leica Biosystems Newcastle Ltd., Newcastle upon Tyne, UK) served for hematoxylin counterstaining. The omission of the primary antibodies served as a negative control.

Quantitative immunohistochemistry was accomplished using plane-matched coronal brain sections stained with appropriate antibodies, which produced a brown-colored end-product visible under a bright-field microscope. The coronal brain sections cut at 25 µm intervals throughout the brain per mouse (1.8–1.9 mm posterior to bregma) were used for the analysis.

### Doublecortin labeling and staining

Newly formed neurons were first labeled with doublecortin (DCX), whose expression is specific for newly generated neuronal cells [42]. We utilized the polyclonal antibody GTX134052 (GeneTex, Irvine, California, USA) diluted at 1:500, to detect doublecortin protein, and quantified the number of DCX-positive neurons and the level of their stain intensity with Zen 2.5 software. The density of neural progenitors (DCX+ cells) in the dentate gyrus was calculated in a circle with a diameter of 385 µm^2^ and presented as the number of DCX+ cells per square mm. DCX+ objects with the surface area greater than 10 µm^2^ were taken into account.

### Polysialylated neuronal cell adhesion molecule staining

Polysialic acid (PSA) is a homopolymer whose primary carrier in vertebrates is NCAM [43]. Commonly, DCX hippocampal expression is temporally in-frame with PSA-NCAM expression [42]. The molecule is exceedingly expressed in the brain during development; still, in the adult murine brain, newborn granule cells of the dentate gyrus highly express PSA-NCAM as well [44]. Accordingly, PSA-NCAM is a popular marker to study structural plasticity and neurogenesis in mammals.

In order to detect PSA-NCAM, we utilized the monoclonal antibody 12E3 #14-9118-82 (eBioscience™, Thermo Fisher Scientific, Waltham, Massachusetts, USA) diluted at 1:100. We quantified the PSA-NCAM-positive surface area and intensity within the dentate gyrus. Zen 2.5 with a preset threshold was used to measure these parameters in a circle with a diameter of 220 µm^2^.

### Microtubule-associated protein 2 staining

Microtubule-associated protein 2 (MAP2) is the most abundant brain MAP, which is predominantly expressed in dendrites and neuronal cell bodies during neurite outgrowth and dendritic branching [45].

The recombinant monoclonal antibody diluted at 1:20000 #ab183830 (Abcam, Cambridge, UK) was utilized for the detection of MAP2 protein. Bright-field micrographs of double-labeled sections have been used for quantitative analysis of immunohistochemistry. MAP2-positive objects with definite and unambiguous neuronal morphology within a circle with a diameter of 220 µm^2^ were analyzed.

### Imaging and quantification

The brain sections have been viewed under an automated upright slide scanning microscope Axio Scan.Z1 (Zeiss, Oberkochen, Germany). The images were captured with 20×/0.8 and 40×/0.95 objectives at z-planes of 0.5 µm. An Axio Imager 2 Upright ApoTome Microscope was used to capture images with 100×/1.4 oil immersion objective.

Image analysis was carried out using ZEN Blue 2.5 (Zeiss). A fixed background intensity threshold was set for all sections representing a single type of staining. In order to create high resolution data, the image deconvolution technique of entire z-series, with ZEN 2.5, was utilized. A computer-driven analysis was performed at each of the counting frame locations.

The surface of the immunoreactive area, above the preset threshold, was subjected to the analysis. The image densitometry method was applied to quantify the amount of staining in the specimens. The mean stain intensity of the specific channel was measured and presented as the average value for each treatment group.

### Tissue sampling, RNA extraction, reverse transcription, and real-time polymerase chain reaction

Five animals per group were rapidly decapitated with scissors. Their brains were carefully removed, and entire hippocampi were sampled. Total RNA was isolated from the left hippocampi using the RNeasy Mini Kit (Cat. No. 74104, QIAGEN, Hilden, Germany) following the manufacturer’s instructions including DNase treatment. RNA quantification was performed using Qubit™ RNA HS Assay Kit (Cat. No. Q32852, Invitrogen, Carlsbad, California, US). RNA integrity was measured using Agilent 2100 Bioanalyzer System and Agilent RNA 6000 Pico Kit (Cat. No. 5067-1513, Agilent Technologies, Santa Clara, California, US). cDNA was prepared from 200 ng of total RNA using SuperScript® III First-Strand Synthesis System for real-time polymerase chain reaction (RT-PCR) (Cat. No. 18080-051, Invitrogen) following the manufacturer’s instructions. RT-PCR was performed using TaqMan Ccl11: Mm00441238_m1 (Applied Biosystems, Foster City, California, US) probe. For the normalization of CCL11 RNA levels, ACTB endogenous housekeeping gene control ActB: Mm00607939_s1 was used. PCR was set in triplicates following the manufacturer’s instructions (Applied Biosystems, Insert PN 4444602 Rev. C) in a 10 μL volume using a five ng cDNA template. PCR was run, and the data was analyzed in the StepOnePlus system installed with StepOne Software v2.3 (Applied Biosystems, Foster City, California, US). The quantification was performed using the comparative Ct (ΔΔCt) method [46].

### Statistical analysis

Statistical analysis was conducted with GraphPad Prism 8.0 for Windows (GraphPad Software, San Diego, CA, USA). The significance was set at 95% of confidence. The two-way ANOVA test was used to demonstrate whether the genotype, the treatment, or the interaction between both factors have an impact upon the phenotype. The two-tailed Student’s t-test was performed to compare the means of two groups. The Kolmogorov–Smirnov test served to evaluate the normality of the data distribution. All data are presented as mean values. Throughout the text and in plots, the variability is indicated by the standard error of the mean (SEM).

## Results

### 3×Tg mice show a significantly reduced DCX-positive NPC density, compared to WT, which demonstrate a dissimilar phenotype, and is unaffected by arginase inhibition

Typically, in mice, the newborn cells of the SGZ migrate into the granule layer and extend their dendrites into the molecular layer [42]. Immunohistochemical staining for DCX of coronal serial sections through the dentate gyrus efficiently revealed newly generated cells.

DCX is a microtubule-associated phosphoprotein, which efficiently labels late mitotic neuronal precursors and early postmitotic cells [47], and is widely used as a reliable marker for newly-born neurons in the adult hippocampus [48]. DCX-positive cells express other early neuronal antigens but are deficient of antigens specific for glia or apoptotic cells [49]. DCX expression is profound in dendrites; accordingly, newly-born neurons’ absolute number and dendritic growth can be efficiently evaluated with DCX immunostaining technique.

Seven-month-old male WT animals show characteristic patterns of DCX expression (Fig 1a). The observed DCX-positive cells are distributed heterogeneously within different regions of the dentate gyrus and are arranged in clusters (Fig. 1a, b). Remarkably, in the WT, the vast majority of the hippocampal DCX-positive neurons are situated in the SGZ; still, a substantial part of them are visible in the granular layer (Fig. 1a, b insets). These bipolar cells demonstrate extensive dendritic growth into the molecular layer.

**Figure 1.**
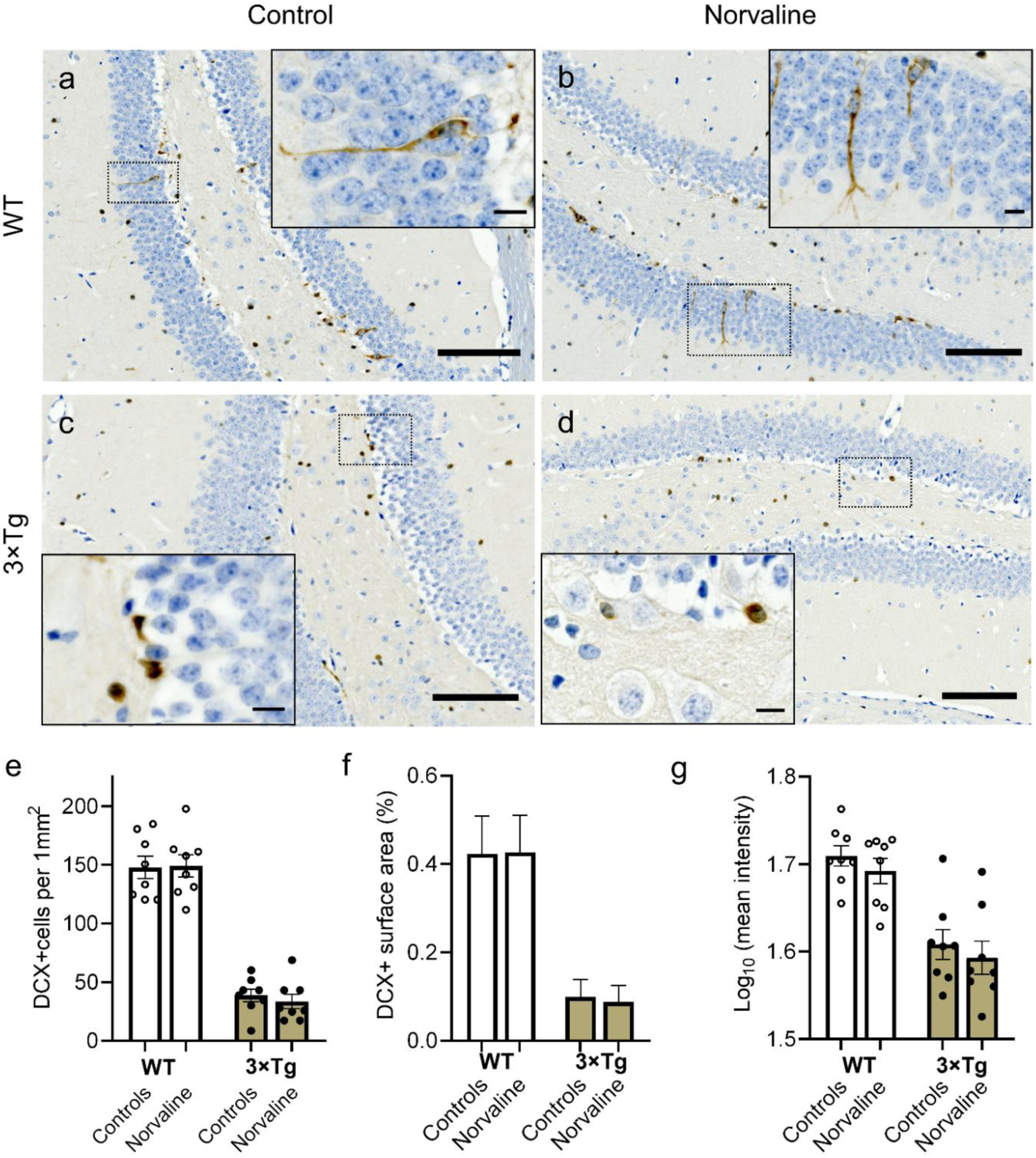
Representative ×40 bright-field micrographs showing the distribution of newly generated DCX-positive cells in the dentate gyrus of 7-month-old male WT (a, b) and 3×Tg (c, d) mice in relation to norvaline treatment. Scale bars represent 100 µm. Brain sections were counterstained with hematoxylin. Newborn cells (brown reaction product) are present in both blades of the dentate gyrus and the hilus area. Insets (a, b) with ×100 magnified views show DCX+ neurons somata in the inner third of the granule cell layer. The cells possess bipolar shape and demonstrate dendritic growth into the outer dentate molecular layer. Insets (c, d) show spherical-shaped and processes-deficient DCX+ neurons clustering in the SGZ. Scale bars indicate 10 µm. The data are presented as means ± SEM, n=8.

In contrast, the 3×Tg mice DCX-positive cells do not exhibit extensive dendrites, and are marginally present in the granular layer (Fig. 1c, d). Two-way ANOVA test reveals a significant effect of genotype on DCX positivity with a significant (p<0.0001; F_1, 28_ = 203.2) reduction in the levels of DCX positive surface area (Fig. 1f), cell density (Fig. 1e), and mean stain intensity (Fig. 1g) in 3×Tg mice as compared to WT age-matched animals. The treatment factor had no significant influence upon these parameters. Additionally, the interaction accounted for less than 0.1% of the total variance.

### Norvaline caused an escalation of the PSA-NCAM levels in the hippocampi of 3×Tg mice, as evidenced by an increase in immunopositive surface area and stain intensity

In order to corroborate the norvaline effects upon the rate of newly generated neurons survival and differentiation rate in adult 3×Tg mice, we tested the hippocampal levels of PSA-NCAM expression via immunohistochemistry. We observed a significant effect of the treatment on PSA-NCAM expression in SGZ, which is characterized by an increase in the levels of stain intensity (Fig. 2d) and the immunopositive surface area (from 0.76±0.2% to 1.86±0.22%) (Fig. 2c). Of note, PSA-NCAM-positive cells are scarcely present in the SGZ of 3×Tg mice and do not penetrate the granular layer (Fig. 2a). In contrast, these neurons are frequent in the SGZ and the granular layer of the 3×Tg mice treated with norvaline (Fig. 2b).

**Figure 2.**
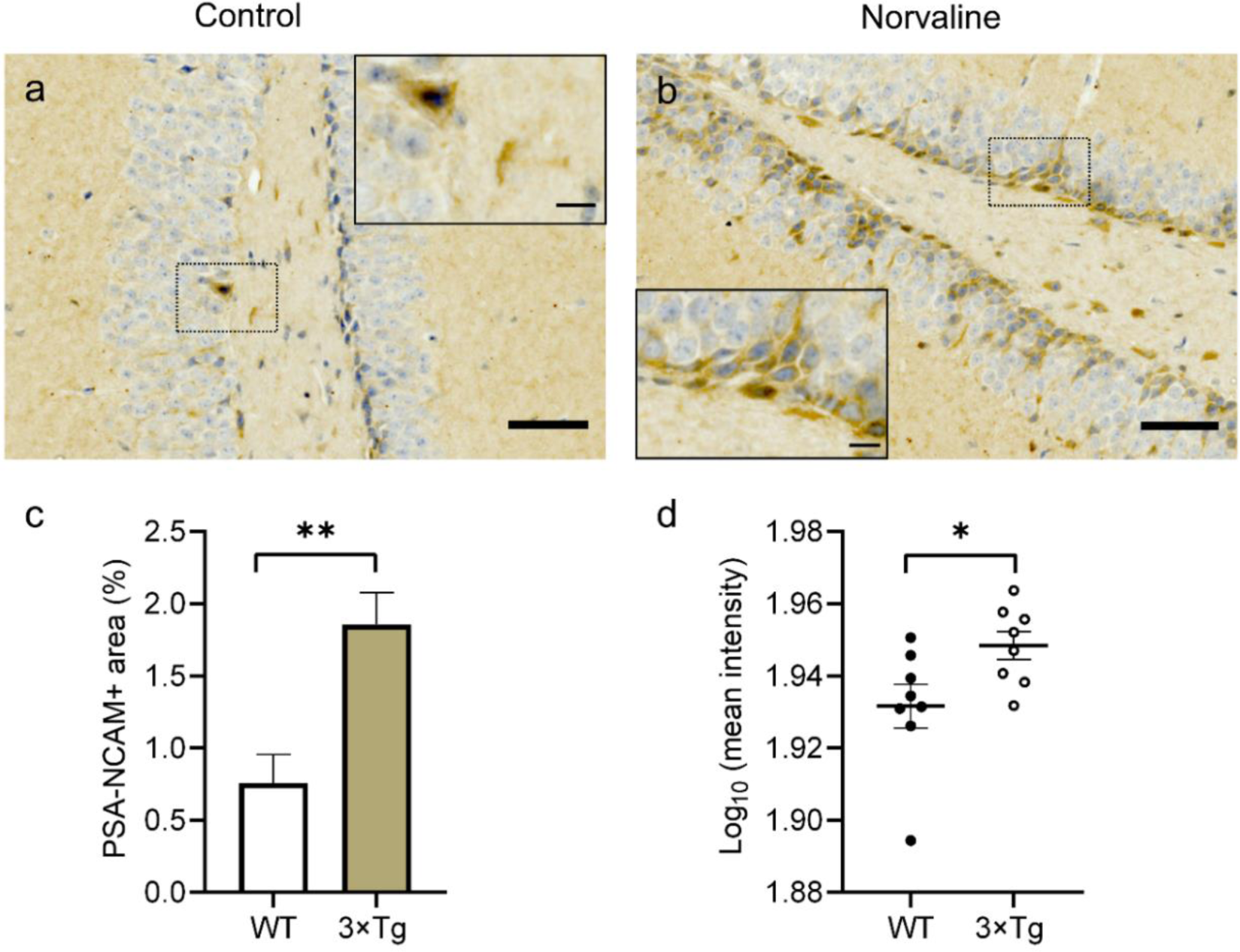
Representative ×40 bright-field micrographs of the hippocampal dentate gyri of 3×Tg mice with ×10 insets (a, b). The SGZ located PSA-NCAM positive cells are marginally present in vehicle-treated animals (a) but show much greater incidence in norvaline-treated mice with penetration into the granule cell layer (b). The treatment is associated with a significant increase in the PSA-NCAM immunopositive area (c) and stain intensity (d). Scale bars 50 µm, insets 10 µm. The data are presented as means ± SEM. *p<0.05, **p<0.01, (two-tailed Student’s t-test), n=8.

### Norvaline rescues neuronal and dendritic loss in 3×Tg mice, as evidenced by MAP2 staining

The dynamic behavior of microtubules is crucial during cell division. MAP2 is a neuron-specific protein stabilizing dendritic microtubules; thus, it serves as a reliable neuronal marker [50]. MAP2-positive neurons possess relatively large cell bodies (more than 20 µm in diameter) and one or more dendrites (50 µm or longer) [51].

We measured the mean stain intensity of the hippocampal MAP2-positive objects and the immunopositive surface area. MAP2-positive objects were quantified in the *cornu ammonis I* (CAI) (Fig. 3e,f) and hilus areas (Fig.3 c,d). Norvaline-treated brains demonstrated robust MAP2 signal, while vehicle-treated brains exhibited a decrement in MAP2 signal as evidenced by two-tailed Student’s t-test. We observed a significant effect of the treatment (p=0.0002, t=4.403, df=22) on MAP2-positive area (with more than three-fold increase) in the CA1 region (Fig. 3h). Stain intensity also demonstrated a significant elevation in CA1 (Fig. 3i). Analysis of the same parameters in the hilus area did not reveal any significant effect, though stain intensity increased with a p-value of 0.059 (Fig. 3g).

**Figure 3.**
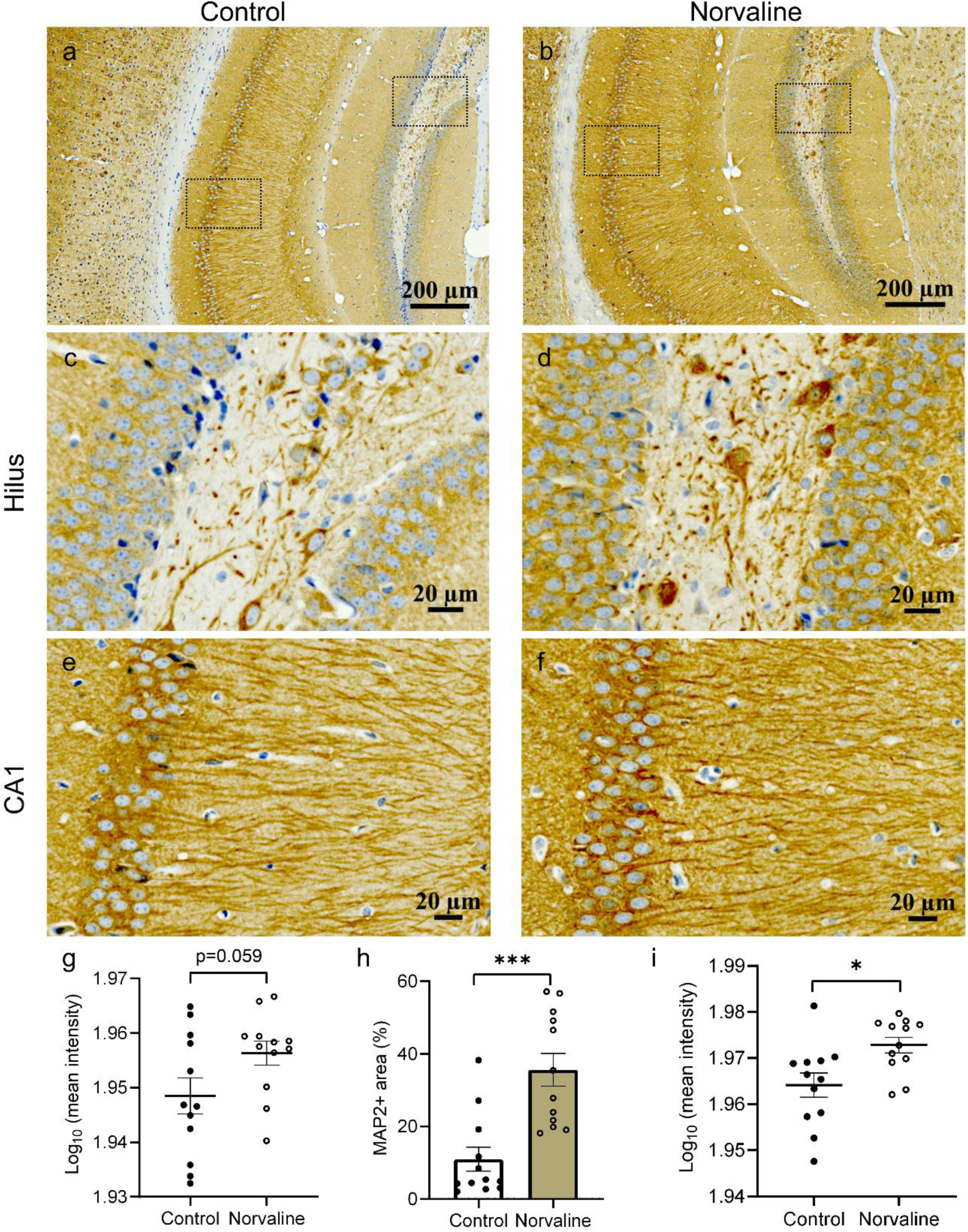
Representative ×20 bright-field micrographs of the 3×Tg mice hippocampi (a,b). ×40 magnification of hilus (c, d) and CA1 region (e, f). Norvaline treatment led to a significant increase in CA1 MAP2-immunopositive surface area (h), and stain intensity (i). The data are presented as means ± SEM (n = 12, four brains per group, three sections per brain). ***p<0.001, *p < 0.05 (two-tailed Student’s t-test).

### Norvaline escalates the transcription levels of CCL11

Eosinophil chemotactic protein (CCL11) has been shown to promote the migration and proliferation of NPCs *in vivo* and *in vitro* [52]. In order to decipher the mechanisms of observed treatment-associated differences in adult neurogenesis, we examined the transcription levels of this β-chemokine in the hippocampi of 3×Tg mice.

Remarkably, the levels of CCL11 mRNA are 79% higher in the norvaline treated mice than in controls. The Student’s t-test demonstrated the significance of the difference between the means of control and treated animals (p=0.0415, t=2.425, df=8) (Fig. 4).

**Figure 4.**
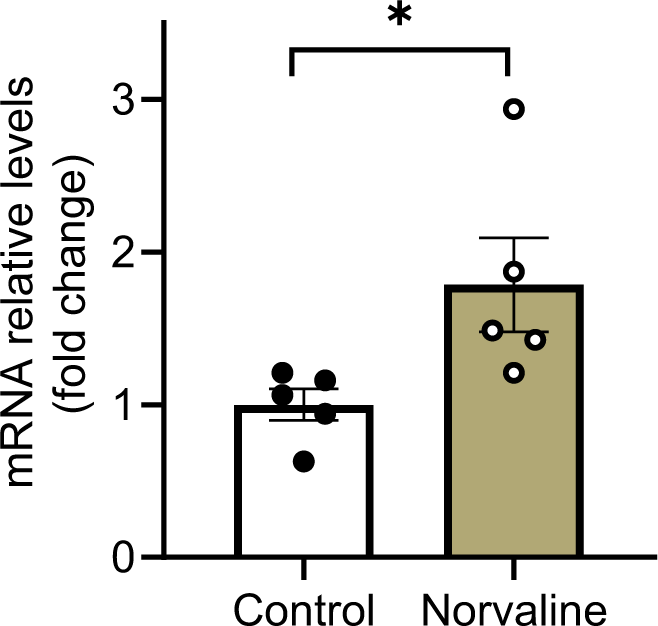
Hippocampal CCL11 mRNA expression levels. RT-PCR analysis of mRNA levels of CCL11 gene. The normalized data are presented as the mean ± SEM (n = 5 brains per group). *p < 0.05 (two-tailed Student’s t-test).

## Discussion

Adult neurogenesis is a complex physiological process that plays a crucial role in the maintenance of normal cognitive functions. This process consists of progenitor cell proliferation, newborn cell migration, and eventually, their maturation [42]. Newborn neurons incorporate into existing functional networks. They are identifiable via various labeling techniques. Dentate gyrus progenitor cells proliferate in the SGZ and migrate into the granular layer of the dentate gyrus. Then, they differentiate and become postmitotic cells with a different phenotype. The new neurons extend their axons to the hippocampal CA3 region and send dendrites to the molecular layer, which functionally integrates them into the hippocampal network [42].

It is worth highlighting that the resident hippocampal precursors are progenitor cells, which are capable of proliferation and multipotential differentiation, though, are incapable of self-renewal [53]. Thus, the extra-hippocampal stem cells generate progenitor cells, which then migrate to the dedicated neurogenic area (SGZ) and proliferate there to produce progeny that differentiate into a population of DCX-expressing newborn neurons. The immature neurons, which express both DCX and PSA-NCAM, decorate the thin lamina underlying the SGZ, and migrate into the granule cell layer [42] (Fig. 1, 2). These cells undergo synaptic integration by sending extensive processes towards the molecular layer and CA3 area, and eventually become typical postmitotic cells.

Adult neurogenesis is an extremely vulnerable process, which is prone to alterations under numerous physiological and pathological conditions. Several lines of evidence suggest a substantial impairment of neurogenesis in AD, which is one of the earliest pathological characteristics of the disease, and its manipulation has been pursued as a potential therapeutic strategy [54].

Various AD animal models show age-dependent neurogenesis deficiency. Decreased proliferation of the hippocampal progenitor cells has been demonstrated in APPswe/PS1dE9 transgenic mice [55]. 3×Tg mice are also characterized by meaningfully impaired adult neurogenesis [3]. Remarkably, various approaches are capable of inducing neurogenesis in adult rodents, including environmental enrichment and enhanced physical activity [56]. Furthermore, numerous studies report a reversal of the decline in neurogenesis in transgenic AD murine models, including 3×Tg mice, as a corollary of different treatment strategies [57-59].

Growing empirical evidence indicates a unique role of NO in adult neurogenesis [60]. Accordingly, several agents have been successfully trialed with a rationale to increase brain NO levels, including arginine [61] and NO-donor supplementation [6]. In this study, we investigated the effects of a different NO-inducing approach upon adult neurogenesis. We utilized an arginase inhibitor, the non-proteinogenic amino acid norvaline, to promote adult neurogenesis in a murine model of AD. We assessed the neurogenesis rate by quantitatively evaluating the proliferation and differentiation of NPCs in the dentate gyrus SGZ. We applied several popular neuronal markers to characterize the different stages of neurogenesis by means of immunohistochemistry.

Previously, we used an advanced proteomics assay to evidence a significant (by 43%) elevation in NCAM protein levels following norvaline treatment in 3×Tg mice brains [15]. In the present study, we scrutinized the spatial patterns of PSA-NCAM hippocampal expression in relation to the treatment. Of note, PSA-NCAM-positive immature neurons have been shown to contribute to the early steps in adult hippocampal neurogenesis, such as proliferation and differentiation [62]. Consequently, PSA-NCAM is used as both a survival and a migration-associated neuronal marker [63]. This biomolecule is required for newly generated neuron survival *in vitro* [64] and *in vivo* [65]. Likewise, the role of PSA-NCAM in migration regulation and in the stimulation of newly generated neuron processes outgrowth has also been suggested [66]. Our methodology revealed a significant increase in the hippocampal granular layer PSA-NCAM positive surface area (Fig. 2 c) and intensity (Fig. 2 d) following the treatment, which implies improvement in the newborn neuron survival rate and accords with our previous results.

It is worth emphasizing that other groups have proven the sensitivity of PSA-NCAM hippocampal levels in AD mice to various treatments and even experiences [67]. In WT animals, PSA-NCAM up-regulation correlates with hippocampal-dependent learning [68]. Therefore, this particular marker reliably indicates the efficacy of the treatment strategy applied and monitors the improvements in hippocampal-dependent function.

In order to evaluate the treatment-associated changes in neuronal and dendritic density, we studied the brain expression patterns of MAP2. MAP2 belongs to a family of heat-stable microtubule-associated proteins, which are responsible for polymerization, stabilization, and dynamics of the microtubule neuronal networks. Accordingly, MAP2 is vital for maintaining neuronal architecture, cell internal organization, cell division, and neuronal morphogenesis [69]. Of note, the levels of MAP2 are significantly diminished in the brains of AD patients [70]. Moreover, *in vitro* studies have shown that Aβ oligomers induce a time-dependent degradation of MAP2 in murine primary cerebral neurons [69]. Another *in vitro* study demonstrated the neuroprotective effect of curcumin, which up-regulates MAP2 expression in human neuroblastoma cells treated with Aβ oligomers [71]. Therefore, MAP2 levels in AD brains are potentially treatment-sensitive and can reflect the treatment efficacy.

It is noteworthy that 3×Tg mice exhibit an early neuronal loss [72] along with a significant reduction in the hippocampal spine density [16]. Previously, we applied Golgi staining and observed a significant increase in hippocampal spine density following norvaline treatment [16]. Here, we assessed the effects of the treatment on the neuronal and dendritic density in 3×Tg mice via quantitative immunohistochemistry with MAP2 antibody. Remarkably, the hippocampi of 3×Tg mice treated with norvaline showed significantly greater MAP2 signal than that of 3×Tg control mice (Fig. 3), These findings point to norvaline rescuing effects on neuronal and dendritic loss, which characterizes the development of memory deficits in 3×Tg mice, and in accord with our previously published data [16].

AD is accompanied by widespread neuroinflammation, and is characterized by chronic microglial activation and overproduction of proinflammatory cytokines [73]. Recent reports have suggested that proinflammatory cytokines, especially tumor necrosis factor-α (TNFα), negatively regulate adult mammal neurogenesis [74], whereas anti-inflammatory cytokines exert the opposite effect [73]. In our previous works, we have shown a significant effect of norvaline treatment upon the rate of microglial activation [16], and the levels of TNFα [15] in the brains of 3×Tg mice. Accordingly, we suggest a supporting effect of norvaline upon adult neurogenesis through the reduction of neuroinflammation.

The small cytokine CCL11, which is produced by neurons in the brain, has been shown to be associated with immune response modulation and protection against neuroinflammation in rats [75]. More recent data strongly implicate the effect of CCL11 on mouse NPCs. Wang *et al*. (2017) utilized a rodent model of hypoxia-ischemia-induced brain damage to demonstrate that CCL11 promotes the migration and proliferation of NPCs [52]. Therefore, we reasoned that norvaline treatment would promote endogenous neurogenesis through neuroprotective factors such as anti-inflammatory cytokines, particularly CCL11. We tested our hypothesis by analyzing CCL11 mRNA expression in the hippocampi of 3×Tg mice in relation to norvaline treatment and found a significant 79% treatment-associated increase. Accordingly, we speculate that CCL11 is at least partially responsible for the phenotype observed in the norvaline-treated mice. This relationship, however, is an assumption until proven by other assays, and further research is needed to shed light on the specific mechanisms of CCL11 induction by norvaline and its role in adult neurogenesis.

Norvaline is an efficient non-competitive arginase inhibitor [76], and this feature, is likely responsible for its neuroprotective properties. Of note, arginine is a common substrate for three enzymes present in several isoforms: arginase, NOS, and arginine decarboxylase (Fig. 5). These enzymes compete for mutual substrate reserves; thus, the overactivation of any one of them leads to the deprivation of others. AD development is associated with arginase overexpression at sites of β-amyloid deposition [12,16], which leads to brain arginine deprivation and also NOS and arginine decarboxylase substrate deficiency. When NOS is deprived of arginine, it undergoes uncoupling, which leads to considerable alterations in its mode of function; these changes reduce the production of NO and generate superoxide anion, which in turn, leads to severe oxidative stress (Fig. 5).

**Figure 5.**
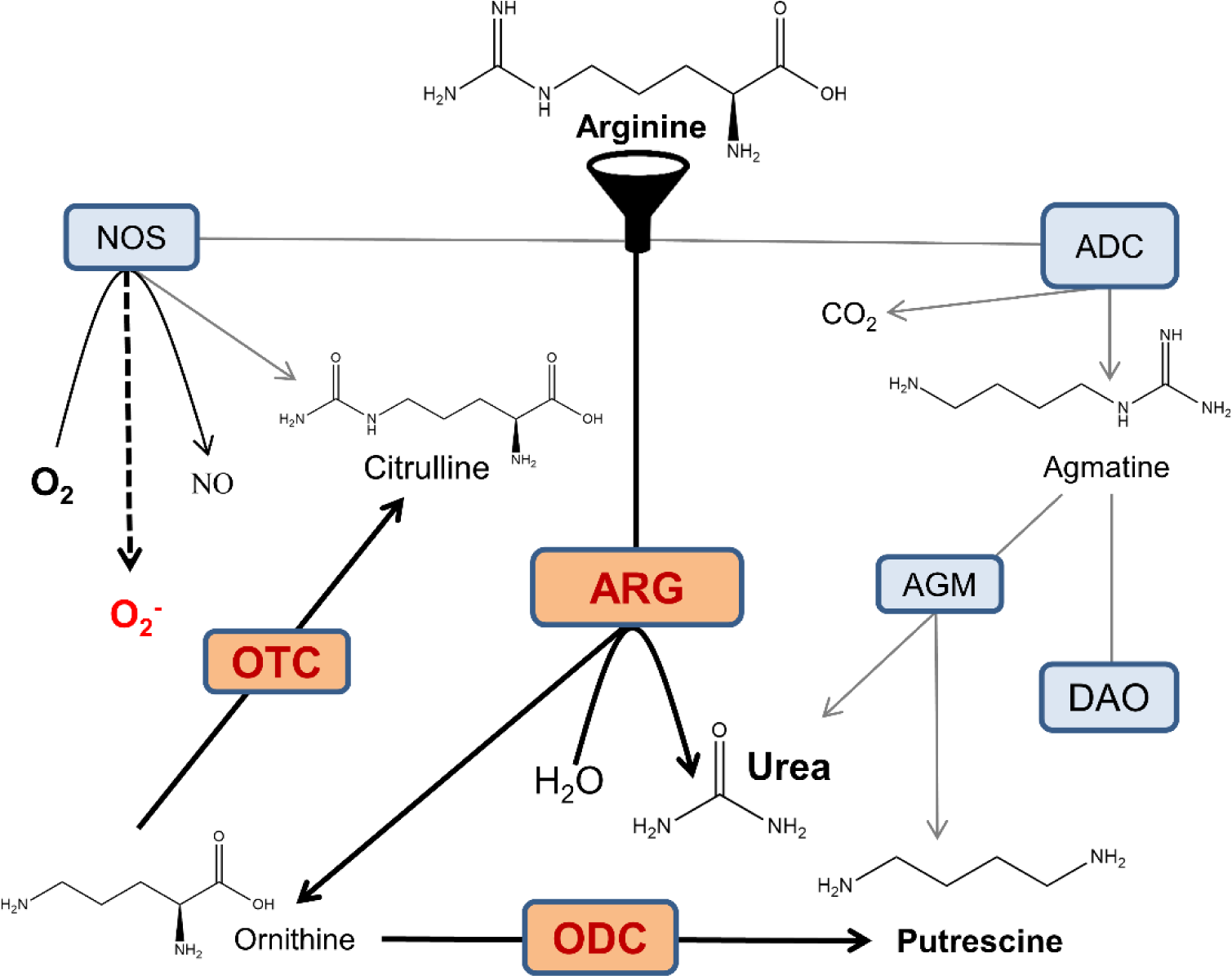
Crossroads of arginine metabolism in the AD brain. NOS oxidatively converts arginine into citrulline and NO. Arginase (ARG) hydrolyzes arginine into ornithine and urea. Arginine decarboxylase (ADC) produces agmatine and carbon dioxide via arginine decarboxylation. Agmatine is utilized in putrescine synthesis via agmatinase (AGM), and alternatively, the neurotransmitter GABA is synthesized in the diamine oxidase (DAO) pathway. Ornithine transcarbamylase (OTC) yields citrulline and phosphate. These pathways interfere with each other via intricate substrate competition mechanisms. For the sake of diagram simplicity, several intermediate steps and byproducts are omitted. In the AD brain, overactive arginase competes with NOS and ADC for the common substrate and reduces the bioavailability of arginine, which limits the production of agmatine and NO, and leads to NOS uncoupling and generation of superoxide anion. Overactivation of ornithine decarboxylase (ODC) leads to a surplus of downstream polyamine products, which can be neurotoxic [77]. Moreover, the gradual oxidation of polyamines by polyamine oxidase is associated with the generation of hydrogen peroxide and leads to oxidative stress [78].

NOS1 has been shown to be chiefly responsible for NO production in the brain and for regulation of vital physiological functions, including neurogenesis [11]. We demonstrated that norvaline upsurges the hippocampal levels of NOS1 [15] and NOS3 [16]; and therefore increases the brain NO content. We speculate that the neuroprotective effects of norvaline are mainly mediated by NO generation and the reduction of oxidative stress, which is a principal characteristic of AD [79].

The role of another arginine-processing enzyme in neurogenesis, arginine decarboxylase, has been recently determined. Arginine decarboxylase is responsible for the conversion of arginine into agmatine [80]. Accordingly, arginase inhibition-associated improvement in the substrate bioavailability leads to an elevation in agmatine brain levels (Fig. 5).

Agmatine has been shown to increase the proliferation rate of cultured hippocampal rat NPCs *in vitro* in a dose-dependent manner and also to induce hippocampal neurogenesis in chronically stressed mice *in vivo* [81]. A more recent study in rats showed that agmatine attenuates traumatic brain injury consequences via promoting neurogenesis and the inhibition of gliosis [82]. Another group proved that agmatine regulates NPCs proliferation and their fate determination in the SVZ [83]. These authors applied immunoblotting and staining to show that agmatine increases MAP2 levels, which supports our findings.

In summary, we have shown that long-term treatment with norvaline promotes NPC survival and differentiation in the hippocampi of 3×Tg mice. Our study offers new insights into the controlling of NPCs function by manipulating the NO microenvironment in the brain. It also provides additional evidence of arginase-targeting benefits in the treatment of AD.

